# DEEP LEARNING ENABLED MULTI-ORGAN SEGMENTATION OF MOUSE EMBRYOS

**DOI:** 10.1101/2022.08.26.505447

**Authors:** S.M. Rolfe, A. M. Maga

## Abstract

The International Mouse Phenotyping Consortium (IMPC) has generated a large repository of 3D imaging data from mouse embryos, providing a rich resource for investigating phenotype/genotype interactions. While the data is freely available, the computing resources and human effort required to segment these images for analysis of individual structures can create a significant hurdle for research. In this paper, we present an open source, deep learning-enabled tool, Mouse Embryo Multi-Organ Segmentation (MEMOS), that estimates a segmentation of 50 anatomical structures with a support for manually reviewing, editing, and analyzing the estimated segmentation in a single application. MEMOS is implemented as an extension on the 3D Slicer platform and is designed to be accessible to researchers without coding experience. We validate the performance of MEMOS-generated segmentations through comparison to state-of-the-art atlas-based segmentation and quantification of previously reported anatomical abnormalities in a CBX4 knockout strain.

**SUMMARY STATEMENT:** We present a new open source, deep learning-enabled tool, Mouse Embryo Multi-Organ Segmentation (MEMOS), to estimate the segmentation of 50 anatomical structures from microCT scans of embryonic mice.

## INTRODUCTION

The IMPC is leading a systematic phenotype evaluation of targeted knockout mouse strains with the goal of providing a comprehensive dataset for exploring genotype/phenotype interactions as part of the Knockout Mouse Phenotyping Program (KOMP2), a National Institute of Health Common Fund project (Dickinson *et al*., 2016). Currently, 7,022 of the 23,000 targeted genes have been evaluated. As part of the standardized phenotyping pipeline, high-resolution 3D micro-CT fetal imaging is collected for sub-viable and lethal strains. So far, 267 lines at E14.5/E15.5 with an average of 6 embryos, have been scanned and new lines are continually added. The high-resolution micro-CT imaging data, along with large sample of wild-type control group provides the capability to detect even subtle differences in organ size and shape in knockout strains (Dickinson *et al*., 2016; Wong *et al*., 2012).

3D imaged fetal specimens are amenable to quantitative phenotyping by measuring the organ sizes and shapes, and their deviation from the normative sample can be a strong indicator of the role a particular knocked-out gene may play in normal development of the organ. However, segmentation of whole-body micro-CT scan images can be a challenging problem due to variation in shape across specimens, the low contrast of soft tissue, and complexity of the three-dimensional organ shape. The current best practice is to have an expert manually add these labels, which is time consuming and prone to variability across time periods, individuals, and datasets. A typical manual segmentation of 50 anatomical structures identified in the E15 fetal atlas can take up to a few hundreds of hours to complete, e.g., Wong *et al*. (2012) cited approximately 400 manhours to derive the E15 mouse fetal atlas segmentation. Thanks to the improved in semi-automated interactive segmentation tools, an expert can probably complete similar tasks a little faster, but in our experience still necessary to commit anywhere from 40-100 hours to fully segment a single E15 mouse scan from the IMPC collection. While this may be feasible for analysis of individuals or small groups, the increasing availability of 3D imaging in big data applications over the last decade has driven a need for automated pipelines that can support increased the throughput necessary to quantify normal variation and detect small-scale differences in phenotype.

Atlas-based image registration has been long used as a method to automate segmentation. In these methods, one (or more) reference images are chosen to serve as an index for a population. Segments or other annotations placed on a reference image are transferred to individual specimen via dense, deformable image registration that provide voxel correspondences between the reference and subject (Park *et al*., 2003; Xu *et al*., 2015). Atlas-based segmentation has now become widely used to improve the both the speed and accuracy of segmentations, including application to the fetal mouse images from the KOMP2 dataset where it was demonstrated to be sensitive enough to identify abnormalities in knockout strains (Horner *et al*., 2021; Van Der Heyden *et al*., 2019; Wong *et al*., 2012). While the gains provided by atlas-based segmentation are significant, these methods are highly computationally intensive and typically require specialized, high-cost compute clusters that take significant time and expertise to maintain, creating another barrier to open data access and repeatable results. They can also show a bias towards normal anatomy and can be sensitive to the selection of the atlas or multiple atlases (Heckemann *et al*., 2006; Van Der Heyden *et al*., 2019).

Recently, deep networks have become a competitive method for automating segmentation in medical and biological imaging applications, with performance exceeding state-of-the-art atlas-based segmentations and have demonstrated advantages in computational requirements (Hatamizadeh, 2022; Schoppe, 2020; Tang, 2021). Deep learning methods also have the potential for performance improvement in regions where anatomy is highly variable and may not be well modelled by a reference segmentation, which is a limitation on atlas-based segmentation. While the training process to generate a segmentation model is computationally intensive, requiring several hours and specialized hardware, once trained, a deployed network can be comparatively lightweight. Inference of an estimated segmentation for a new specimen typically requires a fraction of the computational time and can optionally be run on a standard desktop computer.

In this paper we present Mouse Embryo Multi-Organ Segmentation (MEMOS), a new deep learning-powered tool for segmentation of fetal mice, with the goal of supporting open-access analysis of imaging data collected as part of the IMPC’s KOMP project. MEMOS is provided as a general tool that can be used in a semi-supervised pipeline for fast and highly accurate segmentations or fine-tuned for customized applications.

## RESULTS

### MEMOS training

The pipeline for development of our trained model for segmentation of fetal mouse scans and its deployment in the opensource, user-friendly MEMOS module is summarized in Figure 1. We implemented the deep leaning network for MEMOS using PyTorch and the Medical Open Network for Artificial Intelligence (MONAI) libraries (Diaz-Pinto, 2022). MONAI is an open-source framework for deep learning customized to work with healthcare imaging, including image input and output functions, data processing and image transformation pipelines. Since the application in this work was non-clinical imaging, we trained our model from scratch rather than tuning one of the provided models with initialized weights available as part of the MONAI library. We chose the UNET with transformers, or UNTER, model architecture (Hatamizadeh, 2022). The UNETR is fully implemented in the MONAI library and is currently one of the state-of the-art model for medical image segmentation, demonstrating leading performance on the Multi Atlas Labeling Beyond the Cranial Vault (BTCV) challenge dataset for multi-organ segmentation (Hatamizadeh, 2022).

**Figure 1:**
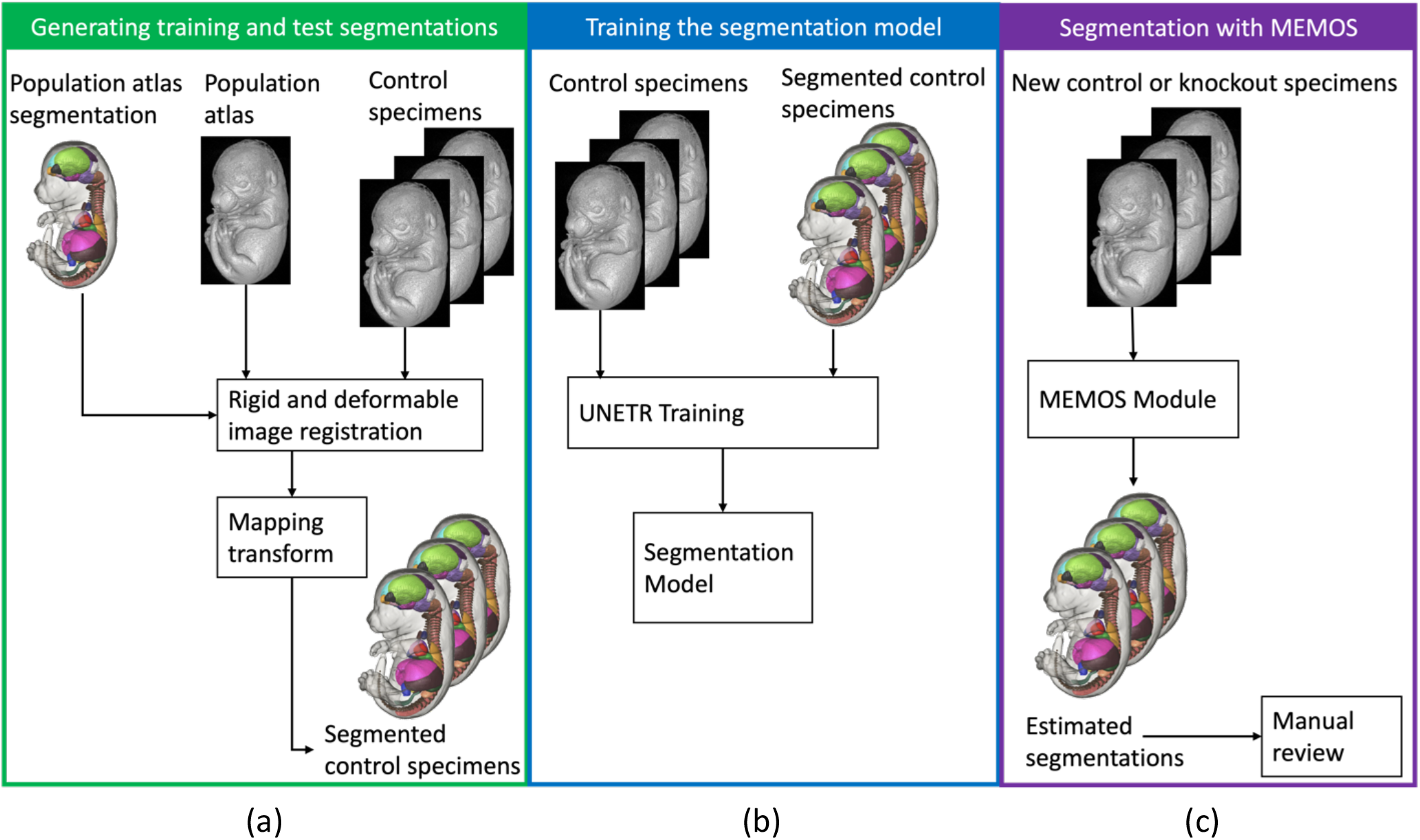
Workflow diagram for training the MEMOS segmentation model and deploying in a user-friendly interface. The three phases of this project included (a) creating labeled training data using atlas-based transfer of the segmentations, (b) training a segmentation model on the atlas-transferred labels, and (c) the deploying the fully trained model in the MEMOS extension to the 3D Slicer application.

### Segmentation Performance

The accuracy of predicted segmentations is evaluated using the Dice coefficient of overlap between the segmentations generated via atlas-based segmentation and those estimated by the MEMOS model. An example of the MEMOS estimated and “ground truth” segmentations for a sagittal slice of a specimen from the validation data set is shown in Figure 2, with similar performance for 24 visible segments. The average Dice score for the fully trained model on the validation dataset is 0.89. To perform an unbiased assessment of the model effectiveness, we evaluated the performance on a test set of 5 specimen withheld from training and validation. For these specimens, the averaged Dice coefficient over all segments is 0.91, indicating similar performance on scans not used to tune the model training parameters. A breakdown of the performance of each segment in the test data set is shown in Table 1. 48 of the 50 segments have a Dice score about 0.8. The vertebrae (Dice score 0.64) and right ventricle of the heart (Dice score 0.63) have a challenging geometry at this image resolution as their width in some dimensions may be as narrow as two voxels. A comparison of the atlas-based and MEMOS segmentations of the right ventricle of the heart for a specimen from the test data set is shown in Figure 3.

**Figure 2:**
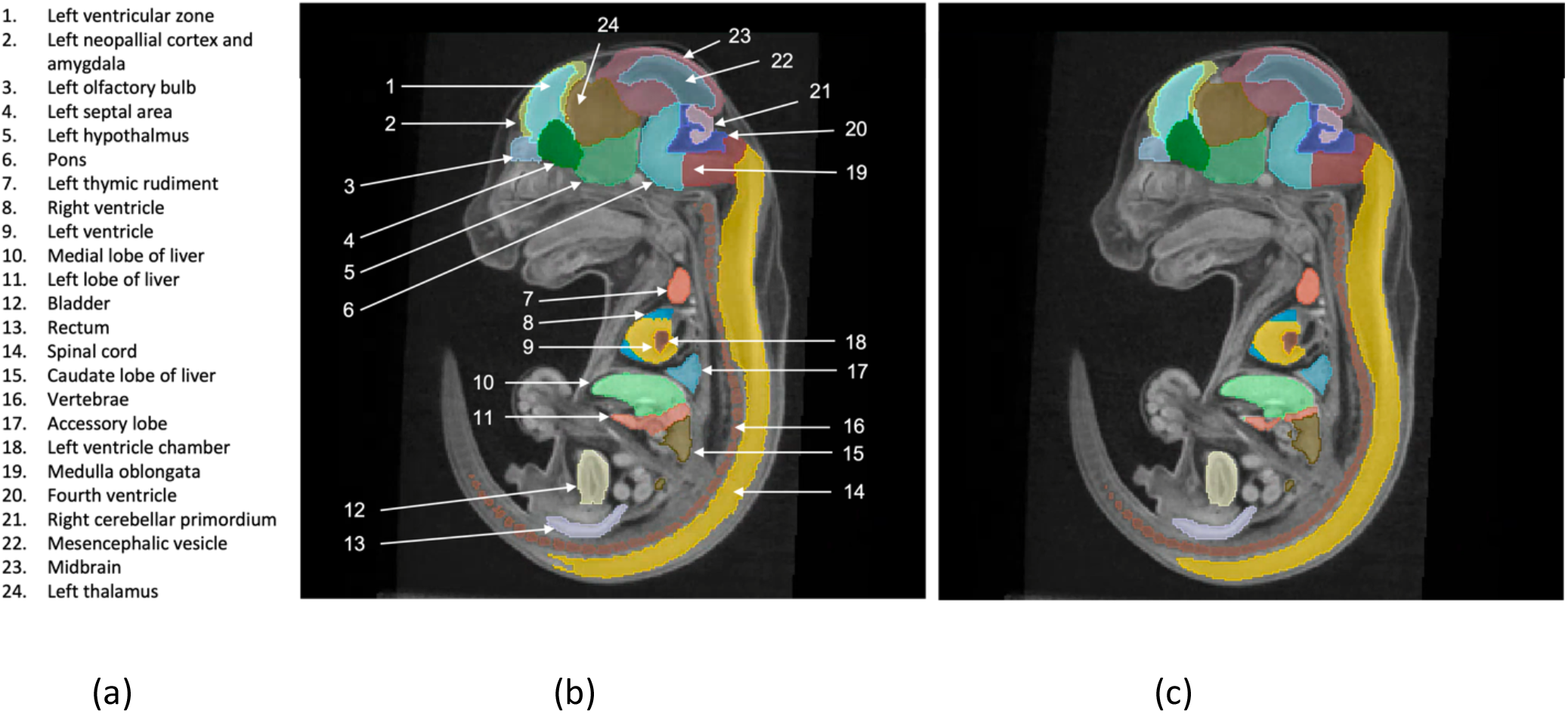
A comparison of the (b) atlas-transferred and (c) MEMOS segmentation labels for a sagittal slice of a specimen from the test data set (not previously seen by the model for training or validation. The 24 labels visible in this figure are defined in (a).

**Table 1.** The average Dice coefficient of overlap between the atlas-based segmentation and the MEMOS segmentation were calculated for 5 specimens withheld from the training and validation data sets. The average Dice coefficient over all segments is 0.91. In this table, the Dice coefficients for all segments is shown, ranked from highest to lowest.

| Segment | Average Dice Coefficient | Segment | Average Dice Coefficient |
| --- | --- | --- | --- |
| left lung | 0.977898 | left thalamus | 0.927241 |
| medial lobe of liver | 0.968217 | midbrain | 0.924903 |
| left lobe of liver | 0.968186 | stomach wall | 0.920919 |
| stomach lumen | 0.966649 | right ventricular zone | 0.920438 |
| left thymic rudiment | 0.966226 | right cerebellar primordium | 0.920331 |
| right thymic rudiment | 0.961125 | rectum | 0.918564 |
| caudal lobe | 0.960622 | caudate lobe of liver | 0.911841 |
| right kidney | 0.956964 | left ventricular zone | 0.905046 |
| right lobe of liver | 0.956306 | left striatum | 0.903752 |
| medulla oblongata | 0.947336 | right striatum | 0.902947 |
| left kidney | 0.947332 | spinal cord | 0.900376 |
| left adrenal | 0.947237 | left cerebellar primordium | 0.898785 |
| cranial lobe | 0.946976 | third ventricle | 0.896352 |
| right hypothalamus | 0.943991 | right ventricle | 0.892408 |
| bladder | 0.942863 | left neopallial cortex and amygdala | 0.890825 |
| pons | 0.942246 | right olfactory bulb | 0.887545 |
| left hypothalamus | 0.938551 | fourth ventricle | 0.884445 |
| right thalamus | 0.937813 | left olfactory bulb | 0.882674 |
| right adrenal | 0.934687 | right neopallial cortex and amygdala | 0.873082 |
| left ventricle | 0.93258 | left ventricle chamber | 0.866304 |
| accessory lobe | 0.929745 | right lateral ventricle | 0.853994 |
| left septal area | 0.929024 | left lateral ventricle | 0.842729 |
| right septal area | 0.928242 | cerebral aqueduct | 0.817713 |
| middle lobe | 0.928217 | vertebrae | 0.641041 |
| mesencephalic vesicle | 0.927903 | right ventricle chamber | 0.637267 |

**Figure 3.**
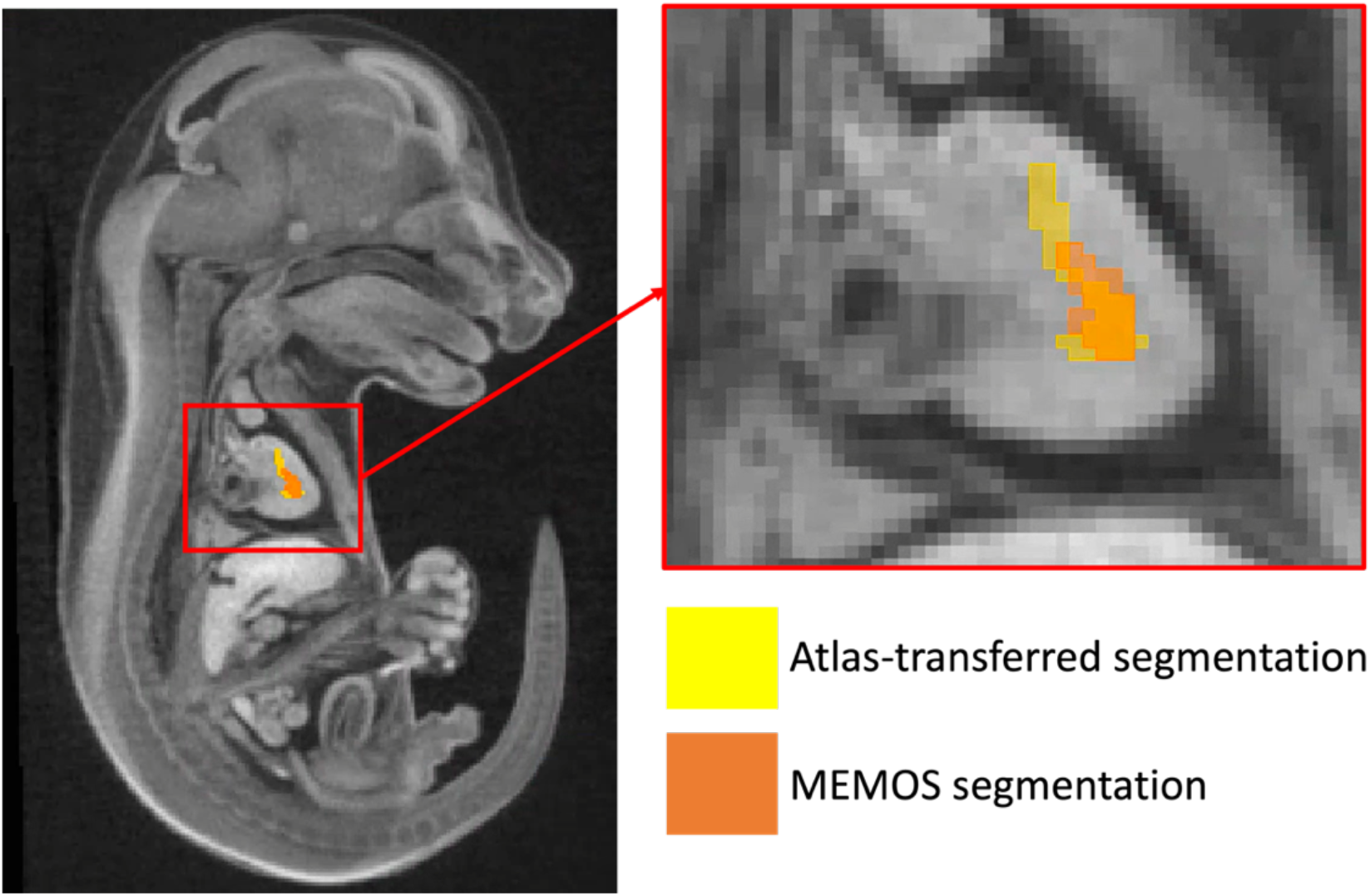
A comparison of the atlas-transferred and MEMOS segmentations of the right ventricle chamber of the heart. The right ventricle chamber of the heart has the lowest performance of the 50 labeled regions, due to its geometry with a width of a few voxels at this image resolution.

### Computational efficiency

To estimate the computational efficiency of the MEMOS module for segmentation, we compared the run time for the volumetric segmentation of 50 labeled anatomical structures for 10 fetal mouse scans not previously seen by the model (used for training or validation) on three different compute systems. The first test system was a Linux server with an AMD Ryzen Threadripper PRO 3995WX processor with 64 cores, 512 GB RAM, and a Nvidia A6000 GPU. The average segmentation time was 21.9 seconds. Compared to the average time for a fully manual segmentation of approximately 40 hours and an atlas-based segmentation on the same high-powered Linux server of approximately 6 hours, this represents speed-up factors of 6,575 and 986 respectively. We tested the same Linux system without using the GPU, resulting in an average segmentation time of 412 seconds, which corresponds to a speed up factor of 350 over the manual segmentation and 52 over the atlas-based segmentation. One important feature of the MEMOS module is that it can be run on a generic desktop computer. Although this is not recommended for the most efficient use of the tool, it increases accessibility by removing the need to purchase specialized hardware. For a standard Windows desktop with 64 GB RAM, Intel Xeon W-2125 processor with 4 cores, the average segmentation time was 1719.9 seconds (28.7 minutes). The atlas-based method was not run on a desktop machine for comparison since the extremely long runtime expected would make this approach impractical. The results from these experiments are summarized in Table 2.

**Table 2:** Summary of the computational systems and average run times used to estimate the efficiency of using the MEMOS module for segmentation.

| System | Compute platform | Memory | Average MEMOS segmentation time (seconds) |
| --- | --- | --- | --- |
| Linux server | Nvidia A6000 GPU,<br>AMD Ryzen Threadripper PRO 3995WX, 64 cores | 512 GB RAM | 21.9 |
| Linux server, CPU only | AMD Ryzen Threadripper PRO 3995WX, 64 cores | 512 GB RAM | 412.0 |
| Windows desktop | Intel Xeon W-2125 | 64 GB RAM | 1719.9 |

### MEMOS module

Our trained segmentation model is deployed in the MEMOS module that is available as an extension of the 3D Slicer platform. 3D Slicer is an opensource software package that leverages powerful image analysis libraries such as ITK, VTK, and an increasing number of state-of-the-art Python packages including the MONAI library to support common image analysis tasks and the creation of custom extensions and workflows (Fedorov, 2012; Kikinis, 2014). The module estimates segmentation of new images in an application providing easy access to tools to review and edit the labels and requiring no programming expertise. The MEMOS module interface is shown in Figure 4. The supplemental material that accompanies the paper provides the necessary steps to install and use the MEMOS module.

**Figure 4:**
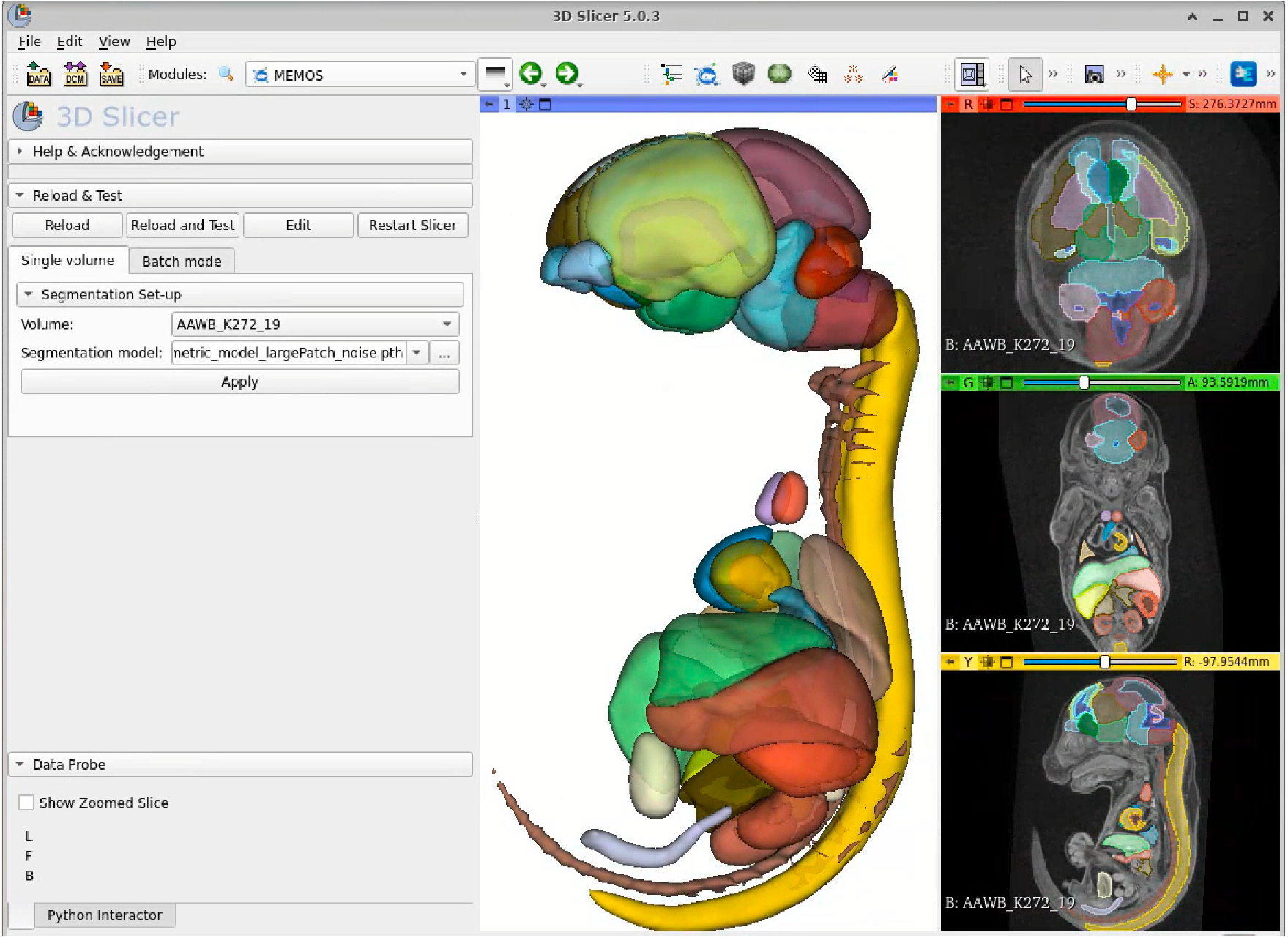
The MEMOS module interface deployed via the 3D Slicer application for segmentation of new specimens. The 3D Slicer application provides interactive visualization, editing, and statistical analysis of the segments.

### Segmentation of knockout mutants with abnormal anatomy

As the primary application of this is segmentation of mouse models to investigate abnormal morphology, the ability to detect small-scale differences in segment size and shape is a key performance metric for MEMOS. To test the segmentation sensitivity, we chose the CBX4 knockout strain from the KOMP dataset, with a phenotype including statistically significant lower volume in the left and right thymic rudiment and the left and right adrenals, as previously reported (Dickinson *et al*., 2016; Liu *et al*., 2014). We used the MEMOS module to segment specimen from the knockout CBX4 strain (n = 8) and compared the adrenal and thymic rudiment volumes, as a percentage of whole-body volume, to those in a test dataset of images not previously seen by the model (n = 11). The average segment volume differentials, detailed in Figure 5, show a statistically significant decrease in volume of the right adrenal and left and right thymic rudiment for the CBX4 knockout strain. The left adrenal also shows a lower average volume, though it does not meet the requirements for significance. An example of the adrenal segmentation of a CBX4 knockout specimen is shown in Figure 6. For the left adrenal, a small region of over-segmentation is visible, likely due to a bias towards normal volume in the training data.

**Figure 5.**
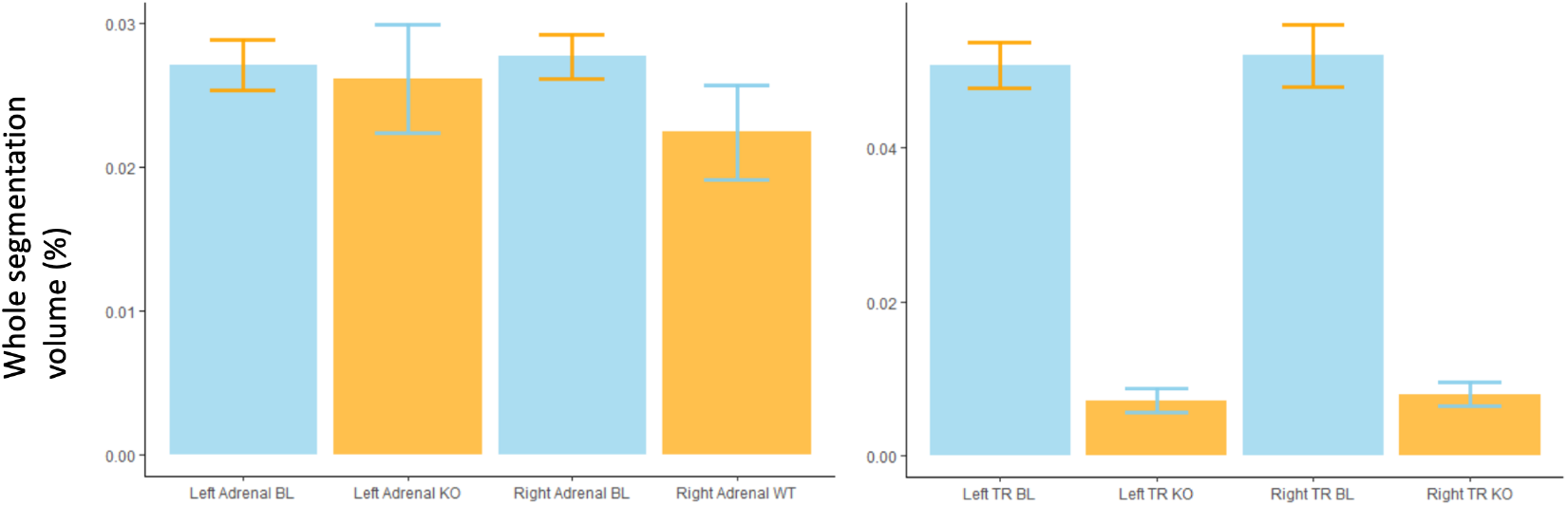
Volumetric comparison of the baseline (BL) and CBX4 knockout (KO) strains for the (a) left and right adrenals and the (b) left and right thymic rudiments. The error bars represent 95% confidence intervals.

**Figure 6.**
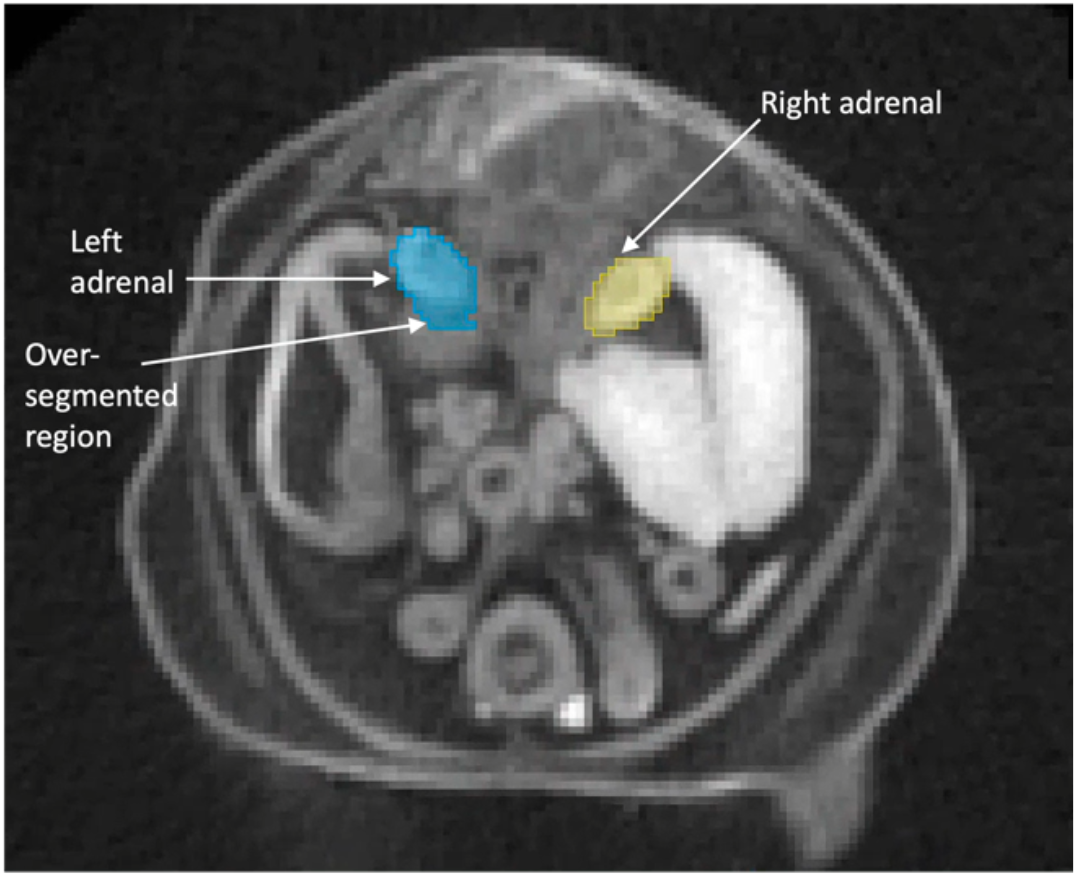
Axial slice from a CBX4 knockout mutant with MEMOS estimated segmentation labels showing over-segmentation of the left adrenal gland.

## DISCUSSION

In this work we have introduced MEMOS, a fully open-source, deep learning-enabled tool for automated estimation of multi-organ segmentations from fetal mouse scans. MEMOS was developed with the goal of supporting segmentation of imaging data collected as part of the NIH Common Fund KOMP2 project. When using a high-performance server, 50 anatomical structures can be estimated within two minutes, although no specialized hardware is required. The accuracy of these segmentations is comparable to gold-standard methods that require multiple orders of magnitude more computation time and significant computational resources. The sensitivity of this tool supports its use with the KOMP2 database on both baseline specimens and strains with known anatomical abnormalities.

The MEMOS segmentations can be generated, reviewed, and if necessary edited within a single open-source application, creating a highly streamlined platform for semi-automated segmentation. The 3D Slicer application provides both the platform for the MEMOS module and an extensive and mature set of interactive segmentation tools that can be applied through the user interface or automatically through simple Python scripts. For example, the MEMOS module can produce small regions of over-segmentation that are spatially disconnected from the remainder of the segmentation. The Segment Editor module within the 3D Slicer application provides an “Islands” tool that identifies the connected components of a segment and can be used to remove small, disconnected regions single button click. The combination of fast initial segmentations produced by MEMOS and accessible, streamlined segment editing tools can be used to quickly generate segmentations for analysis with accuracy that exceeds the MEMOS experimental results.

### Customizing the MEMOS deep learning-model

While goal of this project is to introduce a general-purpose tool for fetal mouse segmentation, the MEMOS deep learning-model can also be customized for specific segmentation tasks. Currently, there are no pre-trained models available for fetal mouse segmentation, so the MEMOS segmentation model provides an important resource for applications where similar datasets are being analyzed. Customizing our model will require some Python programming and access to a deep-learning capable server to retrain the MEMOS model, which can then be loaded and used for segmentation through the existing MEMOS module in 3D Slicer. Here, we briefly address three expected use cases.

### Less-supervised multi-organ segmentation

While the atlas-based registration method for segmentation used to generate our training and validation labels is currently state-of-the-art, it can also be a source of segmentation error. The method tends to be biased towards normal anatomy and atlas selection and is limited in the variation that can be represented. This presents a challenge for segmentating highly deformable internal organs with significant anatomical variation between individuals. While outside the scope of this project, our model could be improved by training on test data that was segmented via the MEMOS model and manually reviewed and if necessary, edited. To further improve the tolerance of MEMOS for high levels of abnormality, such as can be present in the KOMP2 knockout strains, additional examples with abnormality could be included in the training set to model a broader range of local shape variation. Increasing the accuracy of the MEMOS module can reduce the need for manual editing, though a fully automated segmentation will require advances GPU memory available, as discussed in the following section.

### Segmentation of a single or subset of organs

Solving the problem of whole-body segmentation is much more complex than a single segment, or smaller number of segments. For applications where the full set of 50 labels is not necessary, MEMOS could be tuned for segments of interest. To tune MEMOS to a single organ, the model can be further trained on cropped patches from the original, high-resolution volume, centered on that organ. These additional examples can increase the accuracy of the model for that segment. Single-organ networks also provide the benefit of increased efficiency in annotating or editing new training segmentations, since fewer segments need to be placed.

### Segmentation of outside datasets

While MEMOS was trained exclusively on data from the KOMP2 database, our model can be retrained with E15.5 fetal mouse scans from other sources to generate a customized segmentation model for a specific dataset.

### Hardware limitations

Currently the resolution of the MEMOS segmentation network is constrained by the hardware available for deep learning. While state-of-the-art equipment was used to train the UNETR model, the available memory on current GPUs was not sufficient to train on the images at the original resolution. One way to address this constraint is to downsample the images to a resolution that can be directly input into the network. However, the interpolation operations during downsampling can create indistinct boundaries between anatomical regions, breaking biological structure. An alternative strategy which preserves the content of the scans is to break the volume into smaller regions that are used as individual training examples. For applications with a standard space or low variation among subjects, regular patches and their coordinates could be used to preserve positional information lost in sampling. However, full-body segmentation that includes highly plastic abdominal organs have no such expectation of spatial regularity and so we have employed a strategy of sampling randomly selected, overlapping regions. The region size chosen is near the maximum size allowable by our GPU server to preserve spatial context from the volume. In the future, with improved hardware available, the model could be retrained on images at the full resolution to improve segmentation accuracy, towards the goal of fully automated segmentation.

The MEMOS module provides a highly accessible, efficient, semi-automated segmentation pipeline by combining the benefits of a deep learning-enabled segmentation estimates with expert knowledge input via an intuitive interface. MEMOS can produce fast, high-quality segmentations with a minimal amount of user interaction and no requirements for specialized hardware. As the first publicly available pre-trained model for fetal mouse segmentation, we believe the MEMOS deep learning model will be an important contribution to many research applications using fetal mouse imaging.

## MATERIALS AND METHODS

### Imaging Data

As part of the IMPC KOMP2 protocol, iodine contrast-enhanced whole-body micro-CT scans are collected at embryonic stage 15.5 from baseline and knockout strains that are lethal or sub-viable (Dickinson *et al*., 2016). A consensus population average image for stage E15.5 has been calculated from 35 C57BL/6 baseline specimens (Dickinson *et al*., 2016; Wong *et al*., 2012). This average image was manually segmented with 50 labeled anatomical regions in a process taking approximately 400 hours to ensure accuracy (Wong *et al*., 2012).

### Training and validation data

We generated the segmentation model using 91 baseline scans from the KOMP2 dataset. The scans have variable sizes with but have around 250 × 250 × 400 voxels each. In the process of training, the input volumes are randomly sampled with a volume of 128 × 128 × 128 voxels, and the intensity is normalized. The subsampled volumes are augmented by random rotation, affine transformation, intensity shifting, and addition of Gaussian noise.

The training and evaluation strategy was set up to assess how well the MEMOS model could generalize to unseen scans. Towards this goal, we divided the scans into three sets, with 73 images used for training of the model weights, 18 images used to validate the model hyperparameters, and 5 scans withheld for testing. To generate the segmentations required for training and validation, each scan is aligned rigidly and then deformably with the atlas image, in a two-step process implemented using open-source advanced normalization tools (ANT) image quantification library, and its associated R library (ANTsR) (Avants *et al*., 2014; Tustison *et al*., 2021). After the registration, the inverse of the transformation between the atlas and individual is used to map the atlas segmentations into the space of each individual specimen.

### Model architecture

Our model utilizes a UNETR architecture as described in Hatamizadeh et. al. (2022) with encoding and decoding units arranged in a UNET-like contracting-expanding pattern. The 3D input volume is divided into uniform, non-overlapping patches that are flattened into 1D sequences. These sequences are projected via a linear layer into a *K*-dimensional embedding space. A 1-dimensional learnable positional embedding is added to preserve the spatial relationship of the patches. The encoding path consists of a stack of transformers that learn the sequence representations of the input volume. At the bottleneck of the transformer encoder, a deconvolutional layer is used to increase the feature map resolution by a factor of 2. The outputs of the encoding layers are merged with a decoder via skip connections at various resolutions. The decoding units consist of a consecutive 3×3×3 convolution followed by up-sampling through a deconvolutional layer. At the final decoding layer, the output has the original input resolution and is fed into a 1×1×1 convolutional layer that uses a softmax activation function to generate predictions for each voxel in the original volume.

### Model implementation details

Our UNETR model was implemented using the MONAI and Pytorch on a server with 512 GB RAM and A6000 GPUs. The model was trained on the training dataset for 80,000 iterations, using the AdamW optimizer with an initial learning rate of 0.0001. Inference on the validation dataset was reported at increments of 500 iterations. The transformer encoder was implemented with 12 layers, and an embedding dimension size of *K*=768. The non-overlapping patch resolution used was 16. Inference of the segmentation predictions was done using a sliding window approach on the full-resolution testing and validation images. The inference window size is 128×128 x128 with an overlap factor of 0.8 between the windows.

## Supporting information

Supplemental information on installing and running the module described in this work.

## ACKNOWLEDGEMENTS

We would like to thank Dr. Henrik Westerberg for supporting our project and providing advice on accessing the IMPC datasets.

## DATA AVAILABILITY

The baseline fetal mouse imaging data used to train the model are publicly available through the International Mouse Phenotyping Consortium’s website at https://mousephenotyping.org. E15.5 baseline mouse population average and its segmentation were acquired from Toronto Center for Phenogenomics (http://www.mouseimaging.ca/technologies/mouse_atlas/mouse_embryo_atlas.html). The source code for the MEMOS module are available in the GitHub repository: https://github.com/SlicerMorph/MEMOS.

