## Supplemental information on installing and running the module described in this work. for "DEEP LEARNING ENABLED MULTI-ORGAN SEGMENTATION OF MOUSE EMBRYOS"

**Supplementary Information**

**Installing and running the MEMOS module.**

The MEMOS module can be run on Windows, Mac, or Linux OS, on any standard desktop. However, to achieve the maximum benefit to segmentation speed reported in this work, the module should be run on Windows or Linux machine with a CUDA-capable GPU. Installation of the NVIDIA CUDA Toolkit is not covered in this installation guide, please see the CUDA website (https://docs.nvidia.com/cuda/index.html) for detailed for instructions.

1. Download and install 3D Slicer from https://www.slicer.org. Please use the stable release (version 5.0.3) for the most dependable results.
2. Download the MEMOS extension package from https://github.com/SlicerMorph/MEMOS (Click Code button and choose Download zip). Then extract the zip file to a folder.
3. Download the latest trained MEMOS deep learning model from the following links: https://app.box.com/shared/static/4nygg33o70oj5xvnhew11zz5geclus5b.pth

4. **Optional Step for GPU accelerated inference**: To benefit from GPU acceleration provided, your computer needs to be setup with the CUDA development kit. Secondly, this CUDA and this CUDA library needs to match the CUDA version supported by PyTorch (e.g., you can't use CUDA 11.1 with PyTorch, which only supports CUDA 11.3 or 11.6. However, you should be able to use CUDA 11.7 due to backward compatibility). Due to the complexities of this, you need to manually install the Pytorch library for your OS and compute platform inside the Slicer using the following steps. (Note: if you have previously installed another version of the PyTorch library from 3D Slicer, you will need to uninstall it and restart the application before proceeding.)

   First, start the 3D Slicer application and open the Python Interactor.

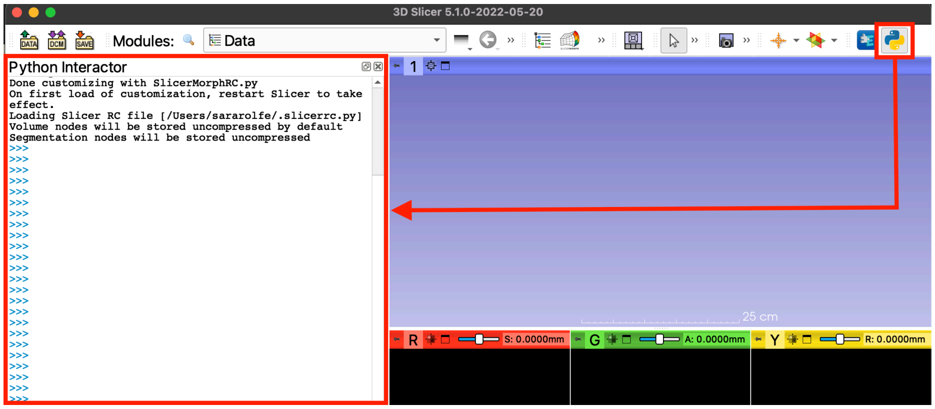


   In the Python Iterator, copy and paste the command from the table below that corresponds your OS and CUDA version. Currently macOS is not supported by the PyTorch pip packages.

| OS | CUDA 10.2 | CUDA 11.3 |
|---|---|---|
| Windows | Not available | pip_install('pip3 install torch torchvision torchaudio --extra-index-url https://download.pytorch.org/whl/cu113') |
| Linux | pip_install('torch torchvision torchaudio') | pip_install('torch torchvision torchaudio --extra-index-url https://download.pytorch.org/whl/cu113') |

For additional details, see https://pytorch.org/get-started/locally/.

5. In Slicer, open the Extensions Wizard module. Choose “Select Extension” from the Extension Tools menu.

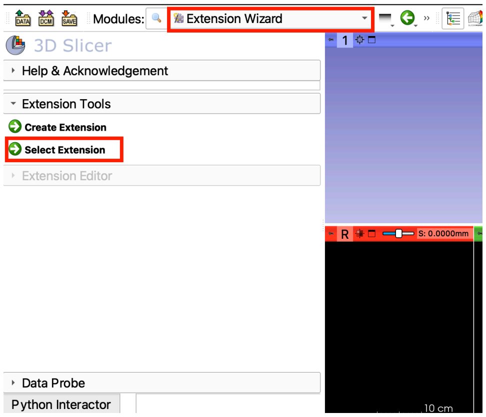


6. Use the file explorer window to select the top-level folder of the MEMOS extension. (note that depending how you extracted the zip archive, this might be one level below the folder you created during extraction.) A pop-up will be displayed asking whether the modules found in the folder should be loaded. Make sure the MEMOS module and option to add to the module search path are checked and click “Yes”.

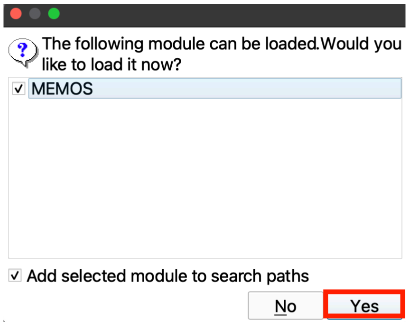


7. Open the MEMOS module. When opening the module for the first time, the application will check for required Python libraries, including Pytorch, MONAI, and their

dependencies. Any libraries not found in the Slicer version of Python will be installed at this time. This step will require an internet connection and may take a couple minutes.

8. The MEMOS module has two modes for segmentation: single volume, where one image is loaded into the Slicer application and segmented; and batch mode, where all images in a folder are segmented and the results are saved to specified output folder.
9. To run MEMOS on a single volume, load the volume into the Slicer application, by dragging and dropping the file into the scene, or using the 3D Slicer "Add Data" dialog box.
10. In the MEMOS module, check that the "Single volume" tab is selected. In the "Volume" field, select the loaded image from the drop-down menu. In the "Segmentation model" field, browse to the MEMOS deep learning model downloaded in step 2. When these fields have been filled in, click "Apply". MEMOS will estimate a segmentation for the volume and once it is complete, it will be displayed in the scene. (Note: If you are not using the GPU acceleration this can take anywhere from 20 minutes to an hour and dependent on the CPU power available in your computer)

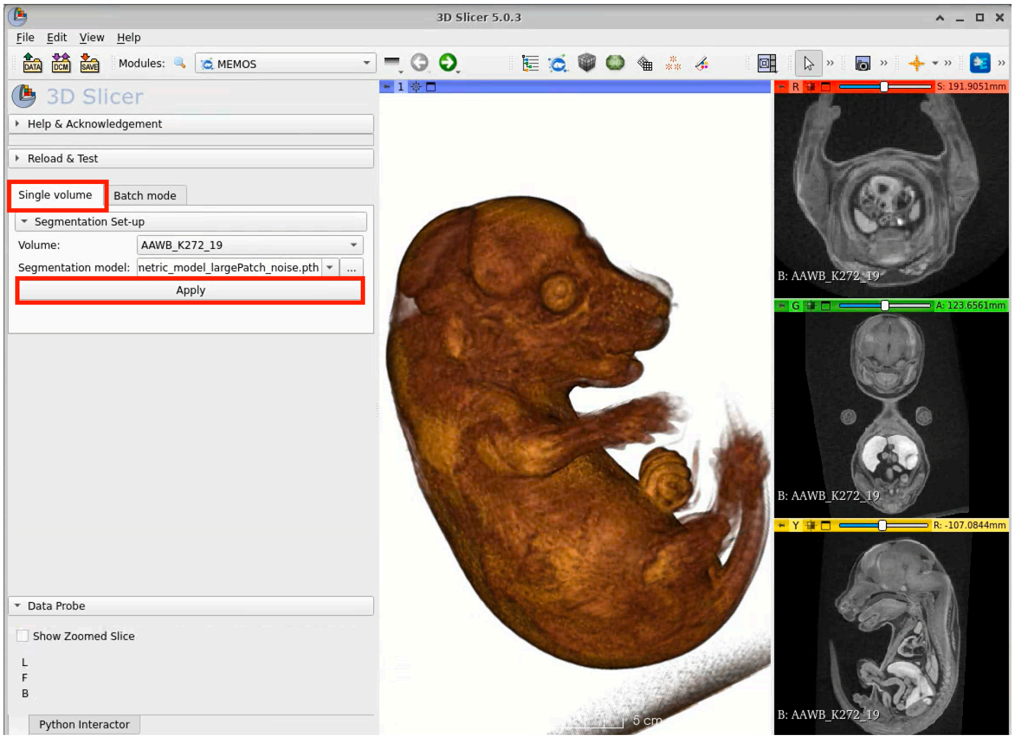


11. To Review and edit the segmentation, browse to the Segment Editor module. For a detailed guide on using the tools in the Segment Editor please see the official user guide: https://slicer.readthedocs.io/en/latest/user_guide/modules/segmenteditor.html and tutorials developed by our research group here: https://github.com/SlicerMorph/Tutorials/blob/main/Segmentation/Segmentation.md

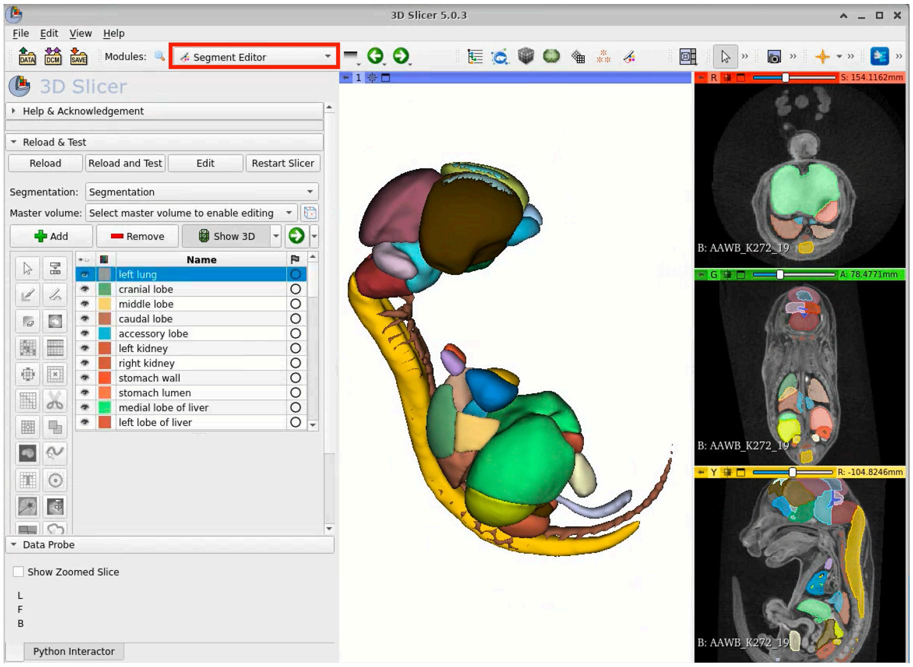


12. Once the segmentation has been reviewed and edited, it can be saved to file using the 3D Slicer “Save Data” dialog box.

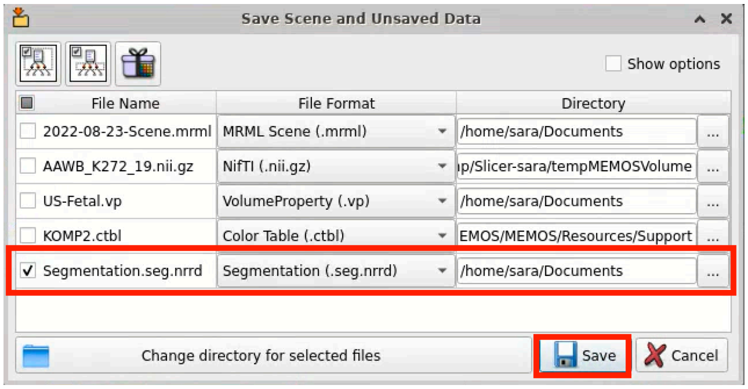


13. To Run MEMOS on each volume in a folder and save the output segmentations, switch to the “Batch mode” tab of the module. Select the folder where the volumes are saved in the “Volume directory” field. In the “Segmentation model” field, browse to the MEMOS deep learning model downloaded in step 2. Select the folder where the segmentations for each volume will be saved in the “Output directory field”. Click “Apply” to generate the segmentations.

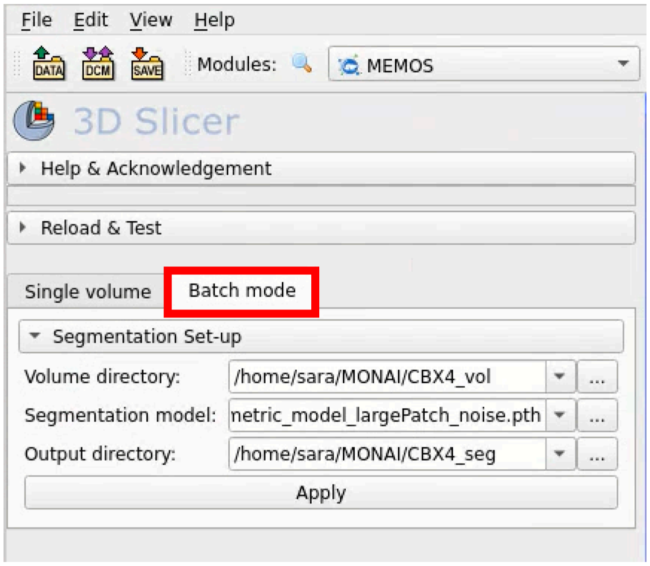


14. To review and edit the segmentations produced in batch mode, load them from the selected output file into the 3D Slicer scene as a segmentation and open the Segmentations Editor module.

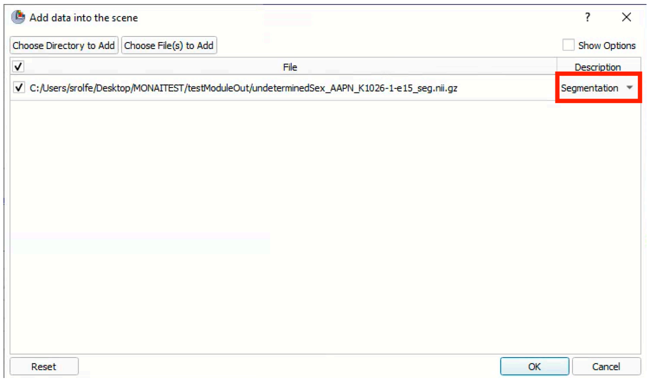
